# IRES-mediated translation of Δ160p53 regulates p53 functions and fine-tunes cancer homeostasis

**DOI:** 10.64898/2026.08.21.744132

**Authors:** Pritam Kumar Ghosh, Piyanki Das, Sahana Ghosh, Risabh Sahu, V Sabari Shree, Subrata Patra, Arindam Maitra, Saumitra Das

**Author notes:** Address for correspondence: Dr. Saumitra Das, Professor, Department of Microbiology and Cell Biology, Indian Institute of Science, Bangalore-560012, India.

## Abstract

Mutations in p53 and its 12 isoforms can alter its functions. As N-terminally truncated isoforms of p53 (Δ40p53, Δ133p53, and Δ160p53) participate in tetramer formation, they are important regulators of cancer fate. Although Δ40p53- and Δ133p53-mediated regulation of cancer is well reported, the mechanism underlying Δ160p53 production and its functional role remains unclear. We investigated the internal ribosomal entry site (IRES)-mediated translation of Δ160p53 and its role in cancer regulation. As differential synthesis of Δ160p53 was observed under different stress conditions, IRES-mediated translation of this isoform was demonstrated using bicistronic luciferase constructs. No cryptic promoters or splicing sites were detected in the IRES sequence. Cell death and late apoptosis were significantly decreased, while proliferation, the number of cells in the S phase, and drug resistance were induced by Δ160p53. Furthermore, Δ160p53 did not induce p53-responsive promoters. RNA sequencing analysis of Δ160p53 overexpression showed similar results, along with the inhibition of other tumor suppressor genes. Overall, our results provide insights into IRES-mediated translation of Δ160p53, which can be considered a novel target for cancer treatment.

## Introductions

The p53 gene (TP53) is 19.2 kb long, spanning over 13 exons (2 alternatively spliced exons and 11 constitutive exons) on chromosome 17 (17p13.1) (Anbarasan & Bourdon, 2019; Joruiz & Bourdon, 2016). p53 is an important nuclear transcription factor and tetrameric protein, which can transactivate several target genes involved in cell cycle regulation and apoptosis (Zilfou & Lowe, 2009). It prevents abnormal cellular proliferation and acts as a tumor suppressor(Finlay *et al*, 1989; Vousden & Prives, 2009). p53 is activated by many stressors, including DNA damage and endoplasmic reticulum (ER) stress (López *et al*, 2015). The p53 protein consists of 393 amino acids and has a molecular weight of 53 kDa. It contains several domains, including two transactivation domains, a proline-rich domain, a DNA-binding domain, a nuclear localization signal, an oligomerization domain, and a C-terminal regulatory domain (Zawacka-Pankau, 2020). TP53 produces 12 isoforms via different mechanisms, including internal ribosome entry sites (IRES), alternative promoters, and alternative splicing (Guo *et al*, 2024; Ray *et al*, 2006). C-terminally truncated isoforms, p53β and p53γ, are produced by the alternative splicing of intron 9 and promote apoptosis (Marcel *et al*, 2014). N-terminally truncated isoforms, namely Δ40p53, Δ133p53, and Δ160p53, can form tetramers with full-length p53 protein (FL-p53) (Zhao *et al*, 2025). Δ40p53 lacks transactivation domain I and acts as a tumor suppressor by inducing apoptosis and cell cycle arrest (Ohki *et al*, 2007; Pal *et al*, 2023; Pal *et al*, 2026; Zhang *et al*, 2022). It is produced by IRES-mediated translation of the mRNA encoding FL-p53 and is regulated by different RNA-binding proteins under stress conditions (Grover *et al*, 2008; Haronikova *et al*, 2019; Khan *et al*, 2015; Sharathchandra *et al*, 2012). The first IRES (IRES1) produces FL-p53, whereas the second IRES (IRES2) produces Δ40p53 (Ray *et al*., 2006). Δ133p53 lacks transactivation domains, a proline-rich domain, and a partial DNA-binding domain (Joruiz *et al*, 2020). Δ133p53 can suppress senescence, cell-cycle arrest, and apoptosis. Furthermore, it can induce proliferation, cellular invasion, and pluripotency in various tissues and cancers (Arsic *et al*, 2015; Campbell *et al*, 2018; Mondal *et al*, 2018; Xie *et al*, 2017). The transcription of Δ133p53-specific mRNA begins from the internal promoter of TP53 at intron 4 (Marcel *et al*, 2010b). Δ133p53 facilitates DNA repair and protects cells from death and senescence in response to DNA damage (Gong *et al*, 2015). Conversely, another N-terminally truncated p53 isoform, Δ160p53, can be produced from the same mRNA variant and does not have transactivation domains, a proline-rich domain, or the first 64 amino acids of the DNA-binding domain (Marcel *et al*, 2010a). p53 mutations can promote the translation of Δ160p53 (Candeias *et al*, 2016). Δ160p53 forms aggregates with FL-p53 in the nucleus and cytoplasm, stabilizes p53, and regulates the transcriptional activity of FL-p53 (Tomas *et al*, 2024; Zhao *et al*., 2025). To the best of our knowledge, the regulation of Δ160p53 synthesis and its function are poorly reported in cancer. Therefore, in this study, we aimed to investigate IRES-mediated translation of Δ160p53 and its role in cancer regulation.

## Results and discussions

### Internal ribosome entry site-mediated translation of Δ160p53

Δ160p53 can be produced from the same mRNA variant from which Δ133p53 is produced (Marcel *et al*., 2010a). The production of Δ160p53 differed in Δ133p53-expressing H1299 cells based on the different stress conditions (Figure 1A). Even though ER stress suppresses global translation, Δ160p53 levels were approximately 28% upregulated after thapsigargin-induced ER stress (Figure 1A). The UNAFOLD algorithm (Markham & Zuker, 2008) predicted a stable secondary structure, consisting of a shorter helix followed by a longer helix interrupted by a bulge. This structure was present within an 81-nucleotide region located immediately upstream of the start codon (AUG) from which translation of Δ160p53 starts (Figure 1B and Supplementary Figure 1A). Thus, these findings predict an IRES sequence (IRES3) immediately upstream of the start codon of Δ160p53. The IRES3 sequence was cloned into a pcDNA construct between Renilla luciferase (R luc) and firefly luciferase (F luc) and was designated pRI3F (pRputativeIRESF) (Figure 1C). The Null bicistronic construct, which lacked an IRES sequence and had previously been characterized (Ray et al., 2006), was used as a negative control. Previously characterized IRES1 and IRES1+2 (containing IRES1 and IRES2 sequences) bicistronic constructs were used to compare the efficiency of IRES3. All bicistronic constructs were transfected into H1299 cells, and normalized luciferase data revealed that IRES3 was as efficient as IRES1 (Figure 1C). To eliminate ribosomal read-through, the putative IRES3 sequence was cloned into a bicistronic construct in which the ΔEMCV sequence was placed upstream of IRES3. The ΔEMCV sequence is a stable RNA structure derived from the IRES of the encephalomyocarditis virus (EMCV) and can inhibit ribosomal read-through (Johannes *et al*, 1999). In the bicistronic construct (peGFPΔEMCVI3Δ160p53 or Δ EMCV Δ160p53), the first gene was eGFP, followed by the ΔEMCV sequence, then the IRES3 sequence, followed by the Δ160p53 coding region. The construct was transfected into H1299 cells, and Δ160p53 was produced along with eGFP (Figure 1D). Internal ribosomal entry was observed after the ΔEMCV sequence, as ΔEMCV can inhibit ribosomal read-through. The same experiment was performed using HeLa cells, and Δ160p53 production was detected in the presence of the upstream ΔEMCV sequence (Supplementary Figure 1B). As the IRES sequence lacks a splice site or cryptic promoter, the presence of an alternative splice site or cryptic promoter was determined in subsequent experiments (Kozak, 2005). To determine the presence of a splice site in IRES3, the pRI3F construct was transfected into H1299 cells, and total RNA was isolated with DNase treatment. Semiquantitative polymerase chain reaction (PCR) was performed using two primer sets targeting Rluc (P3 and P4) and Fluc (P1 and P2). Amplification of the DNA construct (pRI3F) was considered as a positive control in lanes 4 and 9 (Figure 1E). The RNA extract from untransfected cells was considered as a negative control in lane 1. As no band was detected in the absence of reverse transcriptase during the cDNA preparation step (lanes 2 and 5), we confirmed that the total RNA sample was not contaminated with DNA. The specific amplicons (lanes 3 and 6) for R luc and F luc were present as the same size of the amplification bands from the plasmid DNA of pRI3F (lanes 4 and 7), suggesting the absence of alternative splicing site in IRES3 (Figure 1E). To determine the cryptic promoter activity of IRES3, the IRES3 sequence was cloned into a promoter-less vector, pGL-3 basic (I3-pGL3-basic). The normalized luciferase activity and luciferase mRNA levels were not increased in the presence of IRES3 in pGL-3 basic compared with the empty vector (pGL-3 basic) (Figure 1F and G). The PG13-luc construct, which contains a promoter that drives luciferase gene expression, was used as a positive control. These results reveal the absence of any cryptic promoter in IRES3. Thus, the IRES-mediated translation of Δ160p53 was established. To further investigate the potential role of Δ160p53 in cancer progression, we evaluated its effect on cell death and apoptosis.

**Figure 1.**
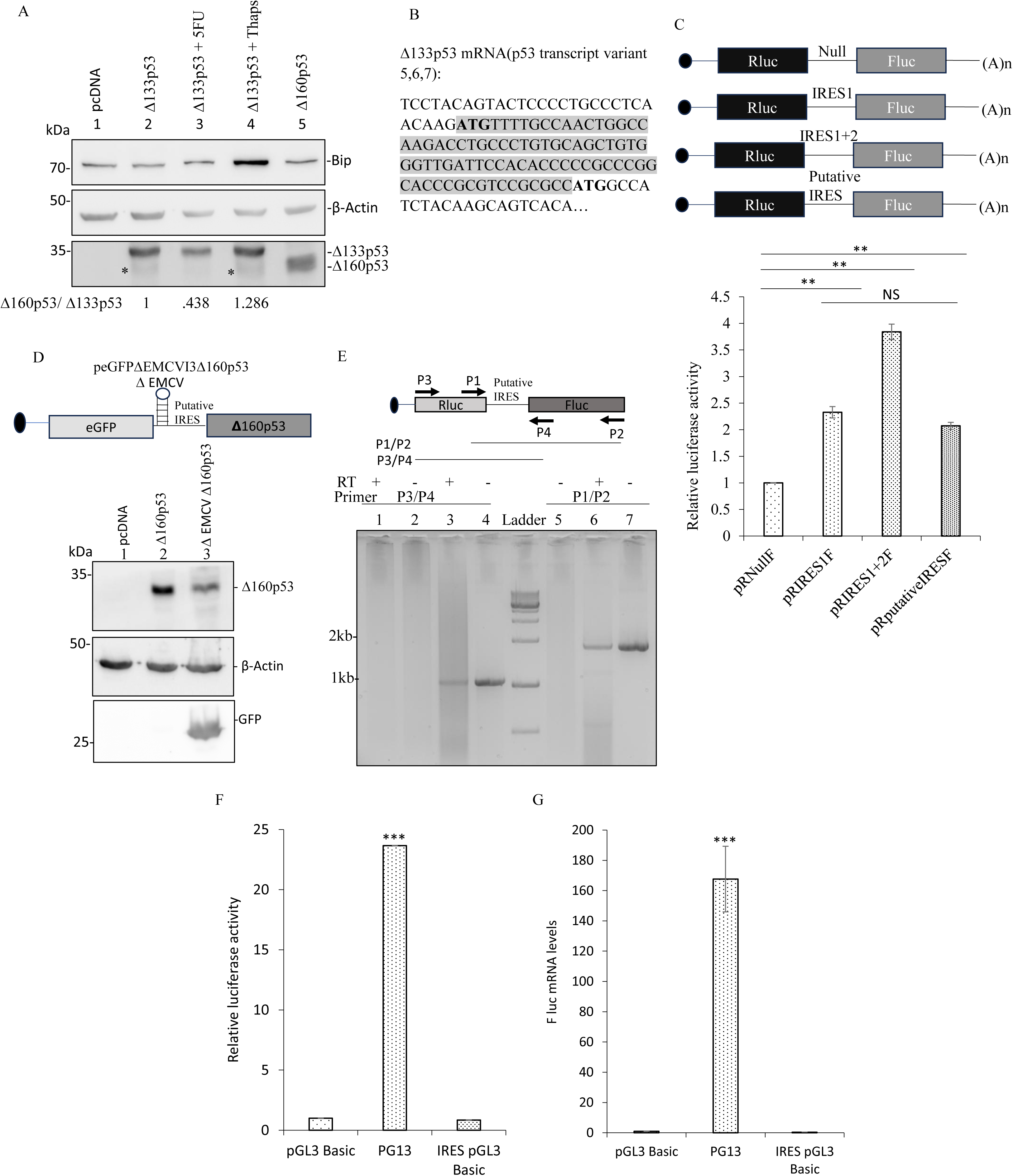
Internal ribosome entry site (IRES)-mediated translation of Δ160p53. (A) Western blot analysis of cell extracts from H1299 transfected with pcDNA and Δ133p53, and treated with 5-fluorouracil (5-FU; 20 µM) and thapsigargin (0.1 µM) for 16 h, probed with BiP antibody, p53 polyclonal antibody, and β-actin antibody. (B) Partial sequence of p53 transcript variants 5,6 and 7, which are known to produce Δ133p53, and the putative IRES sequence was marked. (C) Schematic representation of bicistronic plasmids; H1299 cells were transfected with the plasmids along with pRL-TK. At 48 h post-transfection, relative luciferase activity was calculated in the presence of different IRES sequences and the control sequence (null). (D) Schematic representation of the peGFPΔEMCVI3Δ160p53 construct; western blot analysis of cell extracts from H1299 cells transfected with pcDNA (negative control), Δ160p53 (positive control), and peGFPΔEMCVI3Δ160p53, probed with p53 polyclonal antibody, β-actin antibody, and green fluorescence protein (GFP) antibody. (E) Splicing assay: Semiquantitative polymerase chain reaction (PCR) analysis using two sets of primers (P1/P2 and P3/P4), RNA isolated from pRI3F-transfected (Lanes 2–7) and untransfected (Lane 1) H1299 cells. Lanes 2 and 5 show the reverse transcriptase negative control, and lanes 4 and 7 show the PCR product amplified from the pRI3F bicistronic plasmid (Positive control). (F) Cryptic promoter activity assay: pGL3-basic (Promoter-less luciferase vector as negative control), PG13 (p53 binding sites containing luciferase plasmid as positive control), and IRES-pGL3 plasmids were transfected into H1299 cells along with pRL-TK, and relative luciferase activity was measured. (G) Real-time PCR of Fluc mRNA in H1299 cells transfected with pGL3-basic, PG13, and IRES-pGL3 plasmids. Error bars indicate standard deviation (SD). All experiments were performed in three biological replicates (n = 3). The criterion for significance was determined using a two-tailed Student’s t-test (**P ≤ 0.01 or ***P ≤ 0.001).

### Regulation of cell death and apoptosis by Δ160p53

The pcDNA, Δ160p53, and FL-p53 plasmids were transfected into H1299, and the stable H1299 cell lines were constructed by G418 selection. The Δ160p53 and FL-p53 protein bands confirmed the stable cell lines (Figure 2A). Propidium iodide (PI) staining was performed to detect Δ160p53-mediated cell death. No significant difference was observed in the number of PI-positive cells (dead cells) between Δ160p53 and the vector control in H1299 (Figure 2B and C). FL-p53 can induce cell death, and apoptosis was considered as a positive control (Aubrey *et al*, 2018). FL-p53 considerably induced cell death in H1299 cells, whereas Δ160p53 did not induce cell death (Figure 2C). Different stages of apoptosis were analyzed using annexin V/PI dual staining followed by flow cytometry. We observed that fewer cells undergo late apoptosis upon Δ160p53 induction than upon FL-p53 induction in H1299 cells (Figure 2D and E). In contrast, the number of cells in late apoptosis was the same in Δ160p53 and vector control-containing cells (Figure 2E). The levels of apoptosis markers BAX and PUMA were assessed (Yee *et al*, 2009). PUMA and BAX levels were significantly lower in Δ160p53-containing cells (Figure 2F and G). Therefore, our results suggest that the presence of Δ160p53 suppresses cell death and apoptosis in the lung cancer cell line H1299 compared with FL-p53-containing cells. To further confirm this finding, other cancer-related properties, including cell proliferation, cell cycle progression, and drug resistance, were assessed in subsequent experiments.

**Figure 2.**
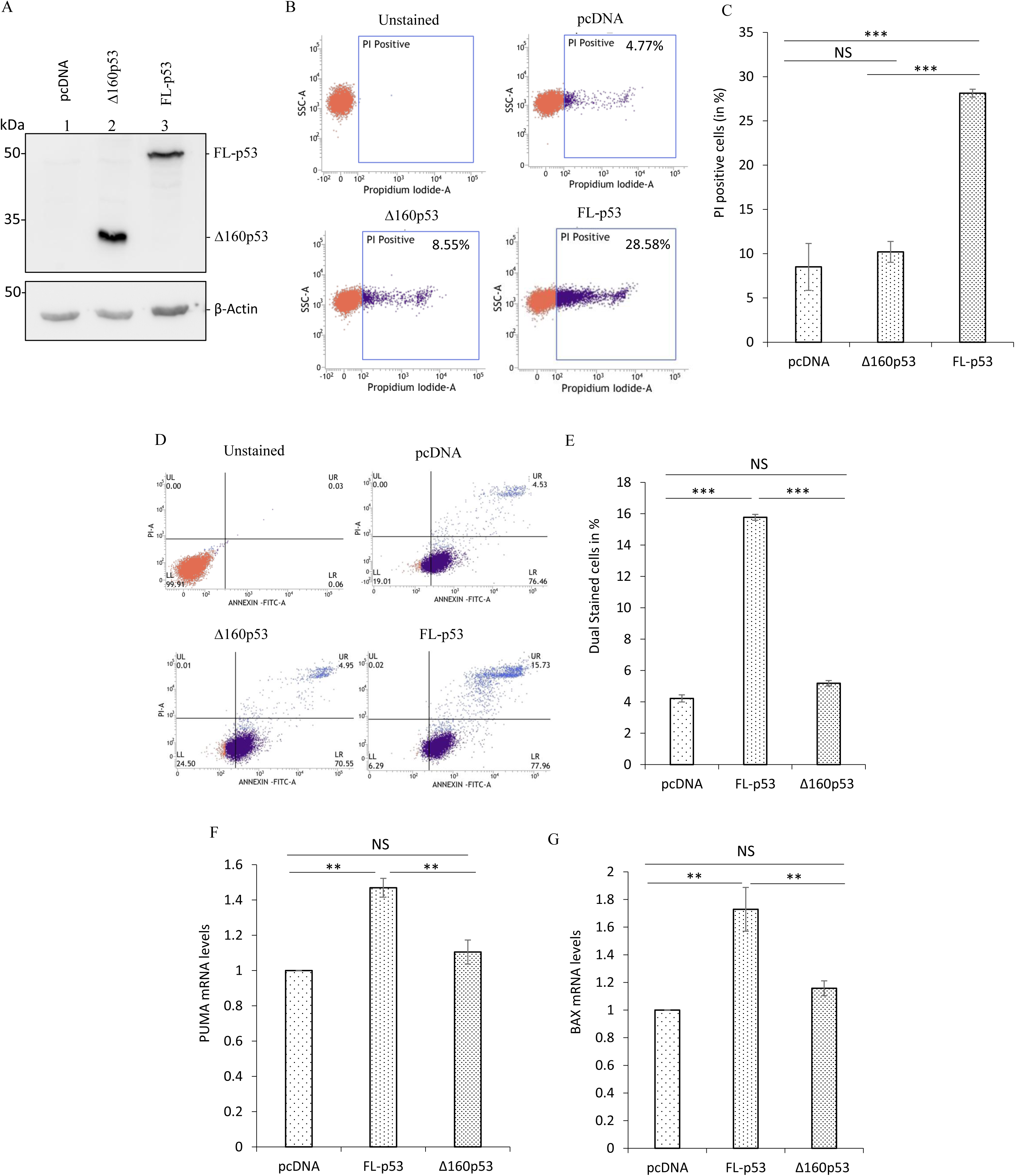
Regulation of cell death and apoptosis by Δ160p53. (A) Western blot analysis of cell extracts from H1299 stable cells expressing pcDNA, Δ160p53, and FL-p53, probed with p53 polyclonal antibody and β-actin antibody. (B) Scatter plot (SSC-A vs. Propidium Iodide-A) for the cell death assay using flow cytometry with propidium iodide (PI) staining of H1299 stable cells expressing pcDNA, Δ160p53, and FL-p53; unstained cells were used to set the gate for PI-positive cells (dead cells). (C) Bar graph analysis of the cell death assay. (D) Scatter plot (PI-A vs. Annexin–FITC-A) for the apoptosis assay using flow cytometry with dual staining of PI and FITC-conjugated Annexin V of H1299 stable cells expressing pcDNA, Δ160p53, and FL-p53. The upper-right (UR) quadrant shows dual-stained cells, which are late apoptotic cells. (E) Bar graph analysis of the apoptosis assay with PI and Annexin V. (F) Real-time PCR of apoptosis marker PUMA in H1299 stable cells expressing pcDNA, Δ160p53, and FL-p53. (G) Real-time PCR of apoptosis marker BAX in H1299 stable cells expressing pcDNA, Δ160p53, and FL-p53. Error bars indicate standard deviation (SD). All experiments were performed in three biological replicates (n = 3). The criterion for significance was determined using a two-tailed Student’s t-test (**P ≤ 0.01 or ***P ≤ 0.001).

### Regulation of cell proliferation and drug resistance by Δ160p53

In pcDNA, Δ160p53, and FL-p53-expressing stable cells, the levels of p53 isoforms were evaluated, and the expression of Δ160p53 and FL-p53 was confirmed (Figure 3A). The WST-1 cell proliferation assay kit was used to determine cell proliferation at different time points under the three conditions. Tetrazolium salt WST-1 was reduced to formazan by cellular dehydrogenases, and the absorbance was detected at 440 nm to determine the viable cell numbers. A standard curve was constructed (Supplementary Figure 2A). No significant difference in cell proliferation was observed at different time points in the presence of Δ160p53 compared with the vector control (Figure 3B and Supplementary Figure 2B). Cell proliferation was significantly downregulated by FL-p53 at 48 and 72 h, as previously reported (Song *et al*, 2015). The number of cells in S phase is a major indicator of cell proliferation (Romar *et al*, 2016). Therefore, we evaluated the differences in the number of cell populations across the different phases of the cell cycle with Δ160p53 and FL-p53. We observed that Δ160p53-containing H1299 cells showed a higher proportion of cells in S-phase than FL-p53-containing H1299 cells (Figure 3C). Our result confirmed that Δ160p53 could not suppress cell proliferation, unlike FL-p53. 5-Fluorouracil (5-FU), a commonly used chemotherapeutic drug, induces p53-mediated cell death and apoptosis (Osaki *et al*, 1997; Pritchard *et al*, 1998; Yang *et al*, 2021). A drug resistance assay was performed using 5-FU at two concentrations (20 μM and 40 μM). The results showed that cell death was not induced by 5-FU treatment in either the vector control or Δ160p53 containing cells at any concentration (Figure 3D and E). Mutated p53 can confer drug resistance in different cancers (Hientz *et al*, 2017). FL-p53 (wild-type)-harbouring cells underwent drug concentration-dependent cell death (Figure 3D and E). Δ160p53 induces drug resistance in H1299 cells. FL-p53 functions as a tumor suppressor by transactivating downstream genes (Hernández Borrero & El-Deiry, 2021). To further elucidate the molecular mechanism underlying the effect of Δ160p53 on cancer cells, including cell death, apoptosis, proliferation, and drug resistance, promoter activation assays and RNA sequencing were performed in the subsequent experiments.

**Figure 3.**
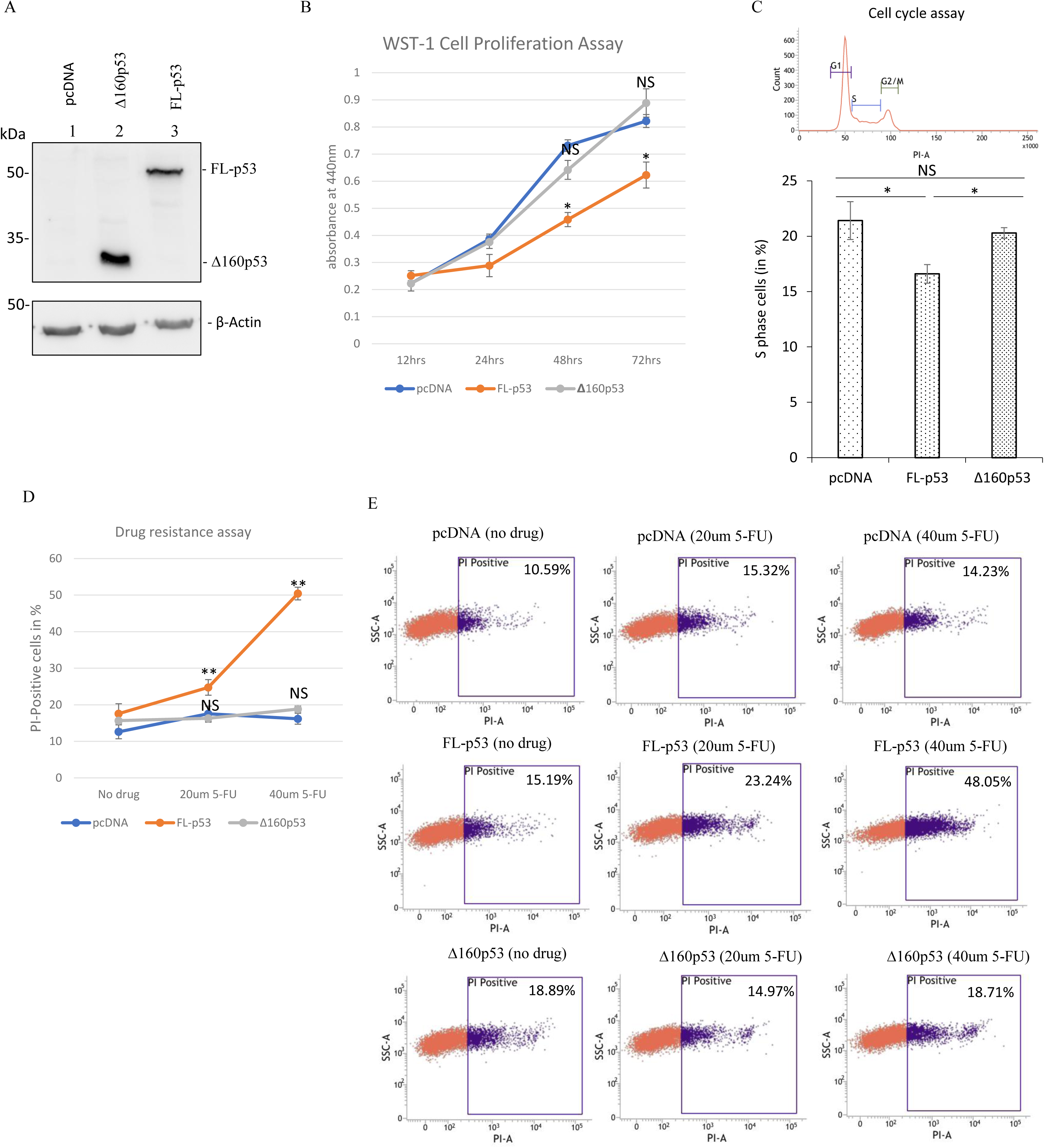
Regulation of cell proliferation and drug resistance by Δ160p53. (A) Western blot analysis of total proteins isolated from pcDNA, Δ160p53, and FL-p53-expressing H1299 stable cells, probed with p53 polyclonal antibody and β-actin antibody. (B) WST-1 cell proliferation assay of H1299 stable cells expressing pcDNA, Δ160p53, and FL-p53 at 440 nm absorbance at different time points (12 h, 24 h, 48 h, and 72 h). (C) Cell cycle assay of H1299 stable cells containing pcDNA, Δ160p53, and FL-p53, and a bar graph of the percentage of S phase in the presence of vector control (pcDNA), Δ160p53, and FL-p53. (D) Drug resistance assay of H1299 stable cells expressing pcDNA, Δ160p53, and FL-p53 treated with 5-FU (20 µM and 40 µM) for 16 h. Cell death was measured using flow cytometry with PI staining. (E) Drug resistance assay: Scatter plot (SSC-A vs. PI-A) for analyzing cell death with PI staining of H1299 stable cells expressing pcDNA, Δ160p53, and FL-p53 treated with 5-FU (20 µM and 40 µM). Error bars indicate standard deviation (SD). All experiments were performed in three biological replicates (n = 3). The criterion for significance was determined using a two-tailed Student’s t-test (*P ≤ 0.05 or **P ≤ 0.01).

### Regulation of promoter activation by Δ160p53

As Δ160p53 is known to impair the DNA-binding activity of FLp53 (Zhao *et al*., 2025), p53 downstream promoter activation was evaluated. The PG13-luc construct contains 13 copies of the FL-p53-binding sequence (el-Deiry *et al*, 1993). Luciferase reporter expression was significantly upregulated in the presence of wild-type p53 (Vilgelm *et al*, 2010). PG13 and pRL-TK were co-transfected with other plasmids (vector control/pcDNA, FL-p53, and Δ160p53) into different cell lines (H1299, A549, HCT116, and HeLa). pRL-TK, which contains R-luc, was used as a transfection control. Our results suggest that Δ160p53 did not induce luciferase activity of PG13. In contrast, FL-p53 can induce the luciferase activity of PG13 to approximately 35-fold compared with vector control in H1299 (Figure 4A). Consequently, the mRNA and protein levels of p21 (a downstream target of FL-p53) exhibited the same trends in H1299 cells (Figure 4B and C). Δ160p53 did not induce p21 expression. Downstream promoter activation was also analyzed in the presence of FL-p53 and the N-terminally truncated isoforms (Δ133p53 and Δ160p53), which mimic the heterotetrameric condition of p53 and its isoforms. FL-p53, mut Δ133p53 (only produces Δ133p53), Δ133p53, and Δ160p53 was overexpressed in H1299 along with PG13 and pRL-TK. The expression of FL-p53 and its isoforms was assessed using western blotting (Supplementary Figure 3A). Under heterotetrameric conditions, both Δ133p53 and Δ160p53 suppressed the FL-p53-mediated induction of luciferase activity (downstream promoter activation). Δ160p53 inhibits more FL-p53 functions compared with Δ133p53 (Supplementary Figure 3B). The effect of Δ160p53 on promoter activation was detected in FL-p53 (Wild-type p53) containing cells (A549, HCT116+/+, and HeLa). Overexpression of Δ160p53 in A549, HCT116+/+, and HeLa cells resulted in a 91%, 84%, and 69% decrease in PG13 luciferase activity, respectively, compared with the vector control (Figure 4D, E, and F). The overexpression of Δ160p53 was confirmed using western blotting (Figure 4G, H, and I). Δ160p53 forms a heterotetramer with FL-p53 (Zhao *et al*., 2025). The results showed that Δ160p53 suppresses FL-p53-mediated transactivation in a heterotetrameric condition when both Δ160p53 and FL-p53 are present. Therefore, Δ160p53 acts as a dominant-negative regulator of full-length p53.

**Figure 4.**
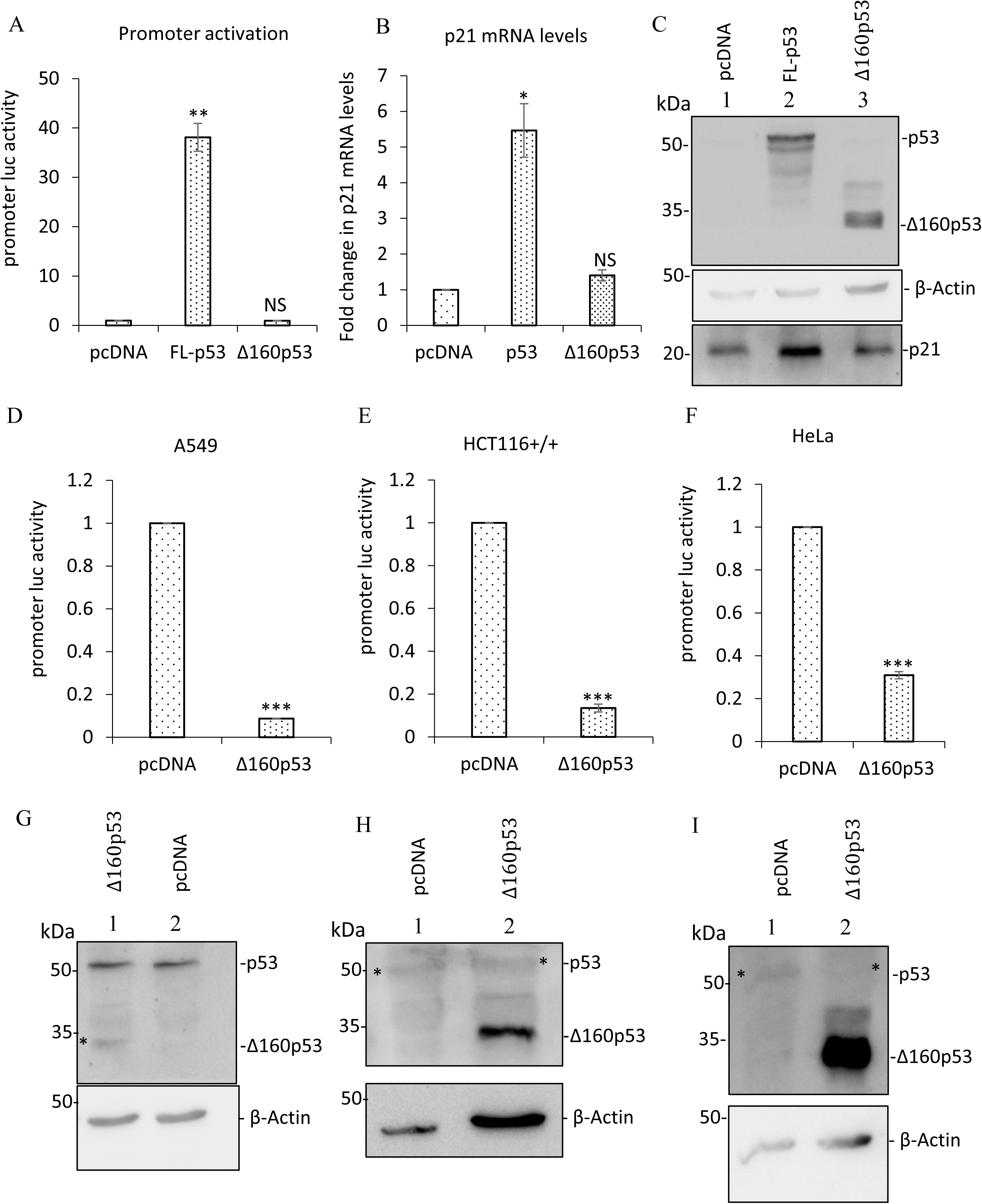
Regulation of promoter activation by Δ160p53. (A) Luciferase assay of H1299 cells after 48 h of transfection with pcDNA, FL-p53, and Δ160p53, and luciferase constructs (PG13 and pRL-TK); the PG13 construct is a p53 binding site-containing firefly luciferase plasmid, and pRL-TK is used as a transfection control. (B) Real-time PCR analysis of p21 mRNA levels in H1299 cells transfected with pcDNA, FL-p53, and Δ160p53 and luciferase constructs (PG13 and pRL-TK). (C) Western blot validation of FL-p53, Δ160p53, and p21 expression from H1299 cells transfected with pcDNA, FL-p53, and Δ160p53 and luciferase constructs (PG13 and pRL-TK), probed with p21 antibody, p53 polyclonal antibody, and β-actin antibody. (D) Luciferase assay of A549 cells transfected with pcDNA, Δ160p53, and luciferase constructs (PG13 and pRL-TK). (E) Luciferase assay of HCT116+/+ cells transfected with pcDNA, Δ160p53, and luciferase constructs (PG13 and pRL-TK). (F) Luciferase assay of HeLa cells after 48 h of transfection with pcDNA, Δ160p53, and luciferase constructs (PG13 and pRL-TK). (G) Western blot validation of FL-p53 and Δ160p53 from A549 cells transfected with pcDNA, Δ160p53, and luciferase constructs (PG13 and pRL-TK), probed with p53 polyclonal antibody and β-actin antibody. (H) Western blot analysis of FL-p53 and Δ160p53 from HCT116+/+ cells transfected with pcDNA, Δ160p53, and luciferase constructs (PG13 and pRL-TK), probed with p53 polyclonal antibody and β-actin antibody. (I) Western blot validation of FL-p53 and Δ160p53 in HeLa after 48 h of transfection with pcDNA, Δ160p53, and luciferase constructs (PG13 and pRL-TK), probed with p53 polyclonal antibody and β-actin antibody. Error bars indicate standard deviation (SD). All experiments were performed in three biological replicates (n = 3). A two-tailed Student’s t-test determined the criterion for significance (*P ≤ 0.05 or **P ≤ 0.01 or ***P ≤ 0.001).

### Molecular mechanism of Δ160p53-mediated regulation of cancer

To identify Δ160p53-regulated factors (genes) that regulate cancer fate, RNA sequencing was performed after transfecting H1299 cells with vector control (pcDNA), FL-p53, and Δ160p53. Expression of FL-p53 and Δ160p53 was identified using western blotting (Figure 5A). RNA sequencing analysis showed that 155 mRNAs were differentially regulated (153 were upregulated, and only two were downregulated) between FL-p53 and Δ160p53 (Figure 5B). No differentially regulated genes were found between pcDNA (vector control) and Δ160p53 conditions in the sequencing analysis, indicating that FL-p53-regulated genes are not regulated by Δ160p53. Pathway analysis was performed using the significantly upregulated mRNAs (≥ 1.5-fold) by FL-p53 (Figure 5C and Supplementary Table 1). Pathway analysis of p53-induced genes identified several downstream tumor suppressor genes associated with cancer-related pathways (Wnt signaling, Hippo signaling, stem cell pluripotency signaling, and pathways in cancer). Genes differentially regulated by FL-p53, ADCY5, Wnt7a, Wnt11, CAMK2B, SFRP4, and CDKN1A, which are known tumor suppressors in different cancers, were chosen for validation and identified by pathway analysis (Figure 5D and Supplementary Table 1) (Bikkavilli *et al*, 2015; Can *et al*, 2024; Jia *et al*, 2022; Shamloo & Usluer, 2019; Wang *et al*, 2018; Wu *et al*, 2021).

**Figure 5.**
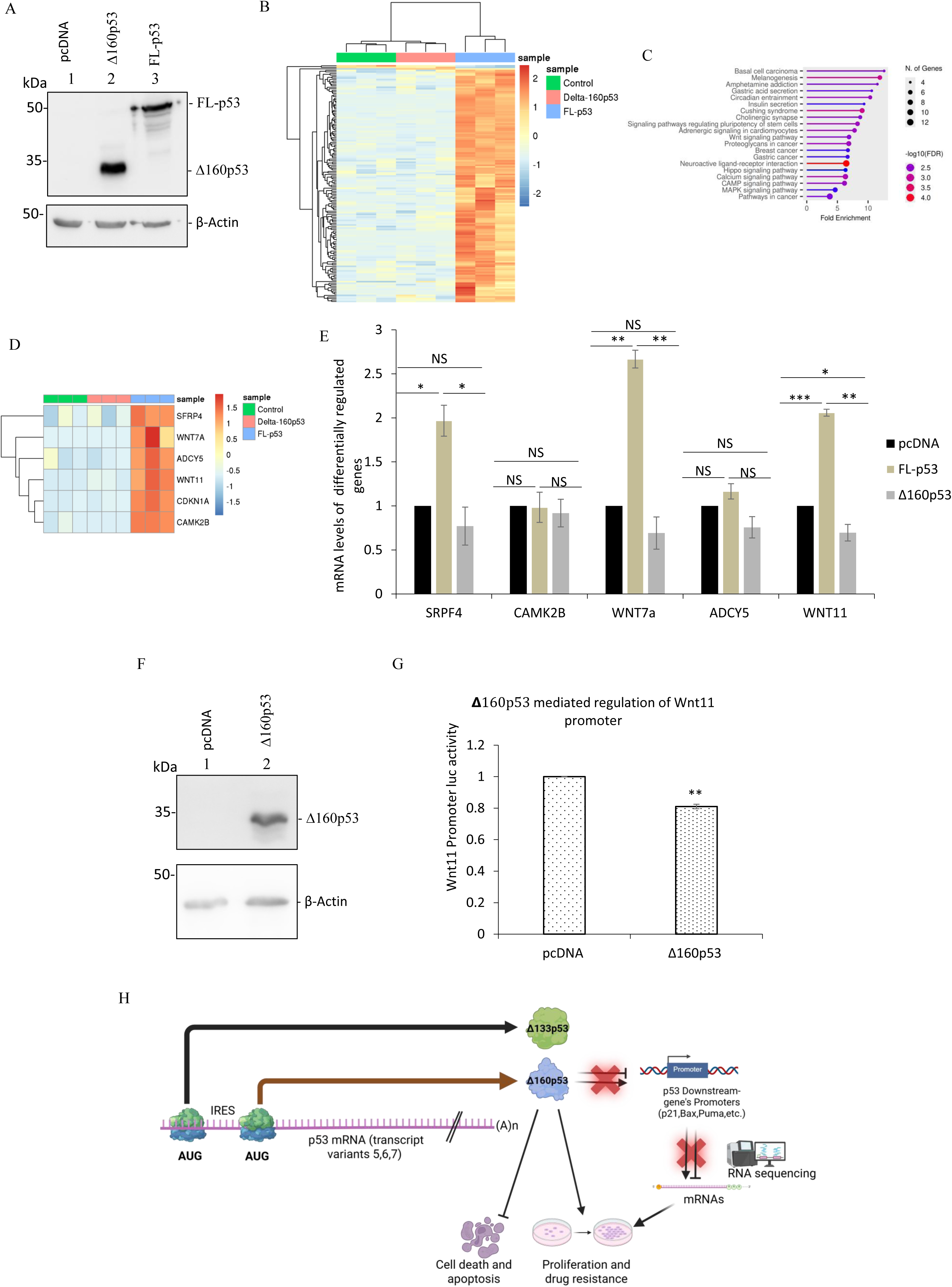
Molecular mechanism of Δ160p53-mediated regulation of cancer. (A) Western blot analysis of total proteins isolated from H1299 cells expressing pcDNA, Δ160p53, and FL-p53H1299, probed with p53 polyclonal antibody and β-actin antibody. (B) Cluster of differentially expressed mRNAs obtained from the total RNA sequencing datasets (heatmap analysis). (C) Pathway analysis of significantly upregulated genes using shinyGO 0.82. (D) Heat map analysis of significantly upregulated cancer-related genes chosen for further validation (p-value < 0.05). (E) Real-time PCR validation of selected genes chosen from RNA sequencing and pathway analysis. (F) Western blot analysis of total proteins isolated from H1299 cells transfected with pcDNA, Δ160p53, and luciferase constructs (Wnt11 promoter luc and pRL-TK), probed with p53 polyclonal antibody and β-Actin antibody. (G) Luciferase assay of H1299 cells after 48 h of transfection with pcDNA, Δ160p53, and luciferase constructs (Wnt11 promoter luc and pRL-TK). (H) Graphical abstract: Δ160p53 can be synthesized from the same transcript variants that are known to produce Δ133p53 and Δ160p53, translated by IRES-mediated translation. The Δ160p53 does not induce cell death or late apoptosis, but it does induce proliferation and drug resistance, as it cannot activate the p53-responsive promoters or their expression at the mRNA level. Error bars indicate standard deviation (SD). All experiments were performed in three biological replicates (n = 3). The criterion for significance was determined using a two-tailed Student’s t-test (*P ≤ 0.05 or **P ≤ 0.01 or ***P ≤ 0.001).

The mRNA levels of SRPF4, Wnt11, Wnt7a, and CDKN1A (p21) were significantly downregulated in the Δ160p53 overexpression condition compared with the FL-p53 in H1299 cells (Figure 5E and 4B). Furthermore, for Wnt11, the decrease was lower than that in the vector control, suggesting that Δ160p53 can independently suppress Wnt11 RNA levels (Figure 5E). To determine the effect of Δ160p53 on Wnt11 promoter activation, the Wnt11 promoter luciferase construct was co-transfected with Δ160p53 in H1299 cells. A marginal but significant decrease in the luciferase levels was observed (Figure 5F and G). Interestingly, Δ160p53 suppressed both Wnt11 promoter activation and its mRNA level (Figure 5E-G). However, Wnt11 levels were not regulated by Δ160p53 in the presence of FL-p53 in A549 cells (Supplementary Figure 4A and B), suggesting that Δ160p53 can regulate Wnt11 expression only under homotetramer condition. As most downstream genes of p53 are not regulated, and Δ160p53 suppresses a tumour suppressor gene (Wnt11), it can neither induce cell death or apoptosis, nor can it suppress proliferation or drug resistance.

In conclusion, this study firmly established IRES-mediated translation of Δ160p53 using the same mRNA that produces Δ133p53. Unlike p53, Δ160p53 does not function as a tumor suppressor. However, it can induce cancer cell proliferation, drug resistance, and suppressed cell death and apoptosis compared with p53 (Figure 5H). Furthermore, heterotetramerization and coaggregation of Δ160p53 with FLp53 can cause a dominant-negative effect (Zhao *et al*., 2025). Consistent with earlier observations, we observed that Δ160p53 cannot regulate p53 downstream genes. However, it can inhibit the functions of FL-p53 and act as a pro-oncogenic factor. Therefore, controlling IRES-mediated translation of Δ160p53 can be a novel therapeutic target for cancer.

## Materials and methods

### Plasmids and constructs

Δ160p53 construct was generated using site-directed mutagenesis of Δ133p53 (a gift from Prof. J.C. Bourdon of the University of Dundee and V Marcel, UK) in the pcDNA construct (ATG to TTG). IRES1 and IRES1+2 bicistronic constructs were used to compare IRESs (Ray *et al*., 2006). The IRES3 sequence was cloned into the HindIII and ECORI sites between R-luc and F-luc in the pcDNA construct (pRI3F). peGFPΔEMCVI3Δ160p53 and I3-pGL3-basic clones were synthesized by Synbio Technologies. The PG13 luc construct was used to determine p53 transactivation. pRL-TK expresses R luc and was used as a transfection control in all luciferase assays. The second AUG of Δ133p53 in pcDNA was mutated (ATG to TTG) via site-directed mutagenesis to generate the mut Δ133p53 construct, which only produces Δ133p53. The pGL4.17 WNT11 (−1025/+46) promoter construct (a gift from Prof. Soon Young Shin of Konkuk University) was used to detect the activation of the Wnt11 promoter by Δ160p53.

### Cell lines and transfections

H1299, A549, HCT116+/+, and HeLa cells were used in this study. H1299 cells are a p53-null human lung adenocarcinoma cell line. A549 (human lung cancer), HCT116+/+ (human colon carcinoma cell line), and HeLa (human cervical cancer cell line) express wild-type p53. All cells were mycoplasma-negative. All four cell lines were grown in Dulbecco’s Modified Eagle Medium (DMEM; Sigma-Aldrich) supplemented with 10% fetal bovine serum (FBS) (GIBCO, Invitrogen). A monolayer of cells at 60–80% confluency was transfected with plasmid constructs using Lipofectamine 2000 (Invitrogen) in nutrient-deficient Opti-MEM (Invitrogen). After 4–5 h of transfection, the culture medium was replaced with DMEM supplemented with 10% FBS. To construct stable cells with pcDNA, FL-p53, and Δ160p53, 5 μg of each construct was transfected into H1299 cells in 100 mm dishes. Subsequently, cells were treated with G418 (800 μg/mL) for 2–3 weeks until 100% of the untransfected cells died. Stable cells were maintained in G418 (500 μg/mL) with complete DMEM. The drug treatments were performed for 16 h immediately before cell harvesting. Subsequently, 5-FU was dissolved in water and used at concentrations of 20 μM and 40 μM to induce DNA damage and assess drug resistance. Thapsigargin was dissolved in DMSO at 100 nM to induce ER stress.

### Luciferase assay

After 48 h of transfection with different luciferase constructs (PG13, pRL-TK, and pRI3F, etc.) using various cell lines, the cells were washed with phosphate-buffered saline (PBS) and lysed in passive lysis buffer. Subsequently, samples were centrifuged at 10,000× g for 10 min (4 °C). The supernatant was collected into new tubes, and luciferase readings were obtained using the Dual-Luciferase® Reporter Assay System (Promega) in a luminometer. All Fluc values were normalized to Rluc values during the analysis.

### Flowcytometry-based assays

Trypsinization was performed during harvesting. Cells were collected in PBS. The collected cells were washed with PBS and fixed with ethanol. Cell death was analyzed using Propidium Iodide (PI) staining. Fixed cells were processed for cell cycle analysis with RNase A treatment and PI staining (10 μg/mL) after ethanol fixation. Cell death was confirmed by PI staining in the drug resistance assay, in which cells were harvested after drug treatment. To determine apoptosis, the Annexin V Apoptosis Assay kit (G-Biosciences) was used as per the manufacturer’s protocol. All flow cytometry data were acquired using a BD FACSVerse.

### Proliferation assay

Five-thousand stable H1299 cells (pcDNA/FL-p53/Δ160p53) were seeded into 96-well plates. CytoScan™ WST-1 Cell Proliferation Assay kit (G-Biosciences) was used to determine cell proliferation at different time points (12 h, 24 h, 48 h, and 72 h) as per the manufacturer’s protocol. The final product, formazan, was detected using a Tecan plate reader at 440 nm. A standard curve was constructed, and the absorbance values were correlated with the cell numbers (Supplementary Figure 2).

### RNA sequencing

Total RNA was isolated from cells using the Purelink RNA Mini Kit as per the manufacturer’s instructions. The quality of the isolated total RNA was determined using an Agilent RNA ScreenTape on an Agilent 2200 TapeStation system, and quantification was performed using a NanoDrop spectrophotometer (Thermo Scientific) and Qubit (Thermo Scientific). Total RNA (500 ng) was used to remove ribosomal RNA using the KAPA RNA HyperPrep Kit with RiboErase (HMR) according to the manufacturer’s instructions (KR1351-v2.17). rRNA-depleted total RNA was used for library preparation, according to the manufacturer’s instructions. Briefly, the first- and second-strand cDNA were prepared from rRNA-depleted fragmented total RNA. Both ends of cDNA were ligated using adapters. Lastly, libraries were enriched for limited-cycle PCR. The quality of the RNA-Seq libraries was assessed using a high-sensitivity D1000 ScreenTape on an Agilent 2200 TapeStation system, and final library quantification was performed using quantitative real-time PCR (RT-PCR). Paired-end 2 × 100 bp sequencing of these libraries was performed using NovaSeq 6000 (Illumina).

### Whole transcriptome data analysis

The 100-bp paired-end reads were quality-checked using FastQC-0.11.7, and only quality control-passed samples were considered in the downstream analysis. Before sequence alignment, a genome index was generated using the human genome assembly FASTA file (GRCh38.p14.genome.fa) and the gene annotation GTF file (gencode.v49.annotation.gtf). The paired-end reads were aligned to the human reference genome along with transcript information. Lastly, the aligned and sorted BAM files (sorted by chromosomal position) were generated for each sample using the aligner STAR (v 2.6) (Dobin *et al*, 2012). The aligned and sorted BAM files were quantified using HTSeq-count (v0.6.1) (Anders *et al*, 2015) which calculated the total raw read counts for each gene in a specific sample. Furthermore, the raw read counts for all samples were normalized for library size using the trimmed mean of M-values (TMM) and expressed as normalized counts per million (CPM) values using edgeR (Robinson *et al*, 2010). DESeq2 (Love *et al*, 2014) was used on raw read counts to identify differentially expressed genes (DEGs) in case samples as compared with control samples. A gene was considered a DEG if its adjusted p-value was < 0.05 and |log2(fold-change)| ≥ 1. Subsequently, unsupervised hierarchical clustering and heat mapping were performed using the log2(normalized CPM + 1) values for all samples, with Euclidean distance and complete linkage. The pathway analysis was performed using ShinyGO version 0.82.

### RNA isolation and RT-PCR

During harvesting, TRI Reagent® (Sigma) was added to the cells, and total RNA was isolated according to the manufacturer’s protocol. DNase I (Thermo Fisher Scientific) treatment was performed to prevent DNA contamination of the RNA samples. The concentration and purity of the RNA were checked using a NanoDrop™ 2000 (Thermo Fisher Scientific). cDNA was synthesized using specific reverse primers and the RevertAid kit (Thermo Scientific) at 42 °C for 1 h. An equal amount of RNA (1 μg) was used for constructing cDNA. For the splicing assay, semiquantitative PCR was performed using Taq polymerase (G-Biosciences) after cDNA preparation. PCR amplification bands were visualized on a 1.5% agarose gel. PCR-mediated site-directed mutagenesis was done with Phusion high-fidelity DNA polymerase (Thermo Scientific) and DpnI (NEB). RT-time PCR was performed using SYBR Green (Thermo Fisher Scientific) and specific primers (Supplementary Table 2) in a QuantStudio™ Real-Time PCR System (Applied Biosystems). The 2ΔΔCt method was used to analyze fold changes in expression. ΔCt = Ct (target gene) − Ct (endogenous control GAPDH), ΔΔCt = ΔCt (target sample) − ΔCt (control sample), and fold change =2(-ΔΔCt). Melting curve analysis was performed after each RT-PCR cycle.

### Western blot analysis

Cells were lysed with radioimmunoprecipitation assay (RIPA) buffer, and the protein concentration was determined using the Bradford assay (Bio-Rad). An equal amount of protein was loaded onto a 12% sodium dodecyl sulfate–polyacrylamide gel electrophoresis gel along with a protein marker (Abclonal, Cat. No. RM02949P) and transferred to a nitrocellulose membrane (Bio-Rad). To detect p53 isoforms, the blots were probed with a rabbit-raised anti-p53 polyclonal antibody (Santa Cruz sc-6243). The BiP antibody (Abclonal, Cat. No. A4908), rabbit anti-GFP-Tag antibody (Abclonal, Cat. No. AE011), and p21 antibody (Abclonal, Cat. No. A19094) were used to probe Bip, GFP, and p21, respectively. Horseradish peroxidase-conjugated anti-β-actin antibody (Sigma) was used to detect β-actin, a loading control in all experiments. Rabbit anti-GFP-Tag antibody (Abclonal, Cat. No. AE011) and β-actin antibodies (Sigma) were horseradish peroxidase-conjugated, and no secondary antibodies were required for detection. For the p53, BiP, and p21 polyclonal antibodies, horseradish peroxidase-conjugated anti-rabbit IgG (Sigma-Aldrich) was used as the secondary antibody. ECL Substrate (Bio-Rad) was used to develop the blots, and images were captured with a Bio-Rad ChemiDoc.

### Statistical analysis

All experiments were performed in three independent biological replicates (n = 3). The graphs were expressed as mean ± standard deviation. Two-tailed paired Student’s t-tests were performed to determine significance. The following p-values were considered significant: P ≤ 0.05 (*), P ≤ 0.01 (**), or P ≤ 0.001(***).

## Supporting information

nothing

## Data availability statement

The sequencing data files are uploaded to the online biorepository forum with the ENA STUDY ID PRJEB112816.

## Author contributions

PKG and SD: Conception and design of studies analysis, interpretation, and article writing. PKG, PD, RS and SV: performing experiments, interpretation of results and article editing. SG, SP and AM: RNA sequencing, analysis and article editing. SD: Funding acquisition, supervision and project management.

## Disclosure and competing interest statement

The authors declare no conflicts of interest.

## Acknowledgements

We thank Prof. J.C. Bourdon (University of Dundee) and V Marcel for Δ133p53 in pcDNA construct. Prof. Soon Young Shin, of Konkuk University, is acknowledged for the pGL4.17 WNT11 (-1025/+46) promoter construct. We acknowledge the sequencing facilities at NIBMG (Kalyani, India) total RNA sequencing and the SD lab members for helpful discussions. This work was supported by the J.C. Bose grant (SD). DBT is acknowledged for the JRF fellowship to PKG.

## Abbreviation

ER Stress: Endoplasmic Reticulum Stress
IRES: Internal ribosome entry site
FL-p53: Full-length p53 protein
R luc: Renila luciferase
F luc: Firefly luciferase
EMCV: Encephalomyocarditis virus
5-Fu: 5-fluorouracil
PI: Propidium Iodide

## Notes

### Competing Interest Statement

The authors have declared no competing interest.

## Reference

Anbarasan T, Bourdon J-C (2019) The Emerging Landscape of p53 Isoforms in Physiology, Cancer and Degenerative Diseases. 20: 6257

Anders S, Pyl PT, Huber W (2015) HTSeq--a Python framework to work with high-throughput sequencing data. Bioinformatics 31: 166–169

Arsic N, Gadea G, Lagerqvist EL, Busson M, Cahuzac N, Brock C, Hollande F, Gire V, Pannequin J, Roux P (2015) The p53 Isoform Δ133p53β Promotes Cancer Stem Cell Potential. Stem Cell Reports 4: 531–540

Aubrey BJ, Kelly GL, Janic A, Herold MJ, Strasser A (2018) How does p53 induce apoptosis and how does this relate to p53-mediated tumour suppression? Cell Death & Differentiation 25: 104–113

Bikkavilli RK, Avasarala S, Van Scoyk M, Arcaroli J, Brzezinski C, Zhang W, Edwards MG, Rathinam MKK, Zhou T, Tauler J et al (2015) Wnt7a is a novel inducer of β-catenin-independent tumor-suppressive cellular senescence in lung cancer. Oncogene 34: 5317–5328

Campbell H, Fleming N, Roth I, Mehta S, Wiles A, Williams G, Vennin C, Arsic N, Parkin A, Pajic M et al (2018) Δ133p53 isoform promotes tumour invasion and metastasis via interleukin-6 activation of JAK-STAT and RhoA-ROCK signalling. Nature Communications 9: 254

Can W, Yan W, Luo H, Xin Z, Yan L, Deqing L, Honglei T, Xiaoyu L, Jiangdong S, Yue X et al (2024) ADCY5 act as a putative tumor suppressor in glioblastoma: An integrated analysis. Heliyon 10: e37012

Candeias MM, Hagiwara M, Matsuda M (2016) Cancer-specific mutations in p53 induce the translation of Δ160p53 promoting tumorigenesis. The EMBO Reports 17: 1542–1551

Dobin A, Davis CA, Schlesinger F, Drenkow J, Zaleski C, Jha S, Batut P, Chaisson M, Gingeras TR (2012) STAR: ultrafast universal RNA-seq aligner. Bioinformatics 29: 15–21

el-Deiry WS, Tokino T, Velculescu VE, Levy DB, Parsons R, Trent JM, Lin D, Mercer WE, Kinzler KW, Vogelstein B (1993) WAF1, a potential mediator of p53 tumor suppression. Cell 75: 817–825

Finlay CA, Hinds PW, Levine AJ (1989) The p53 proto-oncogene can act as a suppressor of transformation. Cell 57: 1083–1093

Gong L, Gong H, Pan X, Chang C, Ou Z, Ye S, Yin L, Yang L, Tao T, Zhang Z et al (2015) p53 isoform Δ113p53/Δ133p53 promotes DNA double-strand break repair to protect cell from death and senescence in response to DNA damage. Cell research 25: 351–369

Grover R, Ray PS, Das S (2008) Polypyrimidine tract binding protein regulates IRES-mediated translation of p53 isoforms. Cell Cycle 7: 2189–2198

Guo Y, Wu H, Wiesmüller L, Chen M (2024) Canonical and non-canonical functions of p53 isoforms: potentiating the complexity of tumor development and therapy resistance. Cell Death & Disease 15: 412

Haronikova L, Olivares-Illana V, Wang L, Karakostis K, Chen S, Fåhraeus R (2019) The p53 mRNA: an integral part of the cellular stress response. Nucleic Acids Research 47: 3257–3271

Hernández Borrero LJ, El-Deiry WS (2021) Tumor suppressor p53: Biology, signaling pathways, and therapeutic targeting. Biochimica et Biophysica Acta (BBA) - Reviews on Cancer 1876: 188556

Hientz K, Mohr A, Bhakta-Guha D, Efferth T (2017) The role of p53 in cancer drug resistance and targeted chemotherapy. Oncotarget 8: 8921–8946

Jia Q, Liao X, Zhang Y, Xu B, Song Y, Bian G, Fu X (2022) Anti-Tumor Role of CAMK2B in Remodeling the Stromal Microenvironment and Inhibiting Proliferation in Papillary Renal Cell Carcinoma. Frontiers in oncology 12: 740051

Johannes G, Carter MS, Eisen MB, Brown PO, Sarnow P (1999) Identification of eukaryotic mRNAs that are translated at reduced cap binding complex eIF4F concentrations using a cDNA microarray. Proc Natl Acad Sci U S A 96: 13118–13123

Joruiz SM, Beck JA, Horikawa I, Harris CC (2020) The Δ133p53 Isoforms, Tuners of the p53 Pathway. Cancers 12

Joruiz SM, Bourdon JC (2016) p53 Isoforms: Key Regulators of the Cell Fate Decision. Cold Spring Harbor perspectives in medicine 6

Khan D, Katoch A, Das A, Sharathchandra A, Lal R, Roy P, Das S, Chattopadhyay S, Das S (2015) Reversible induction of translational isoforms of p53 in glucose deprivation. Cell Death & Differentiation 22: 1203–1218

Kozak M (2005) A second look at cellular mRNA sequences said to function as internal ribosome entry sites. Nucleic Acids Research 33: 6593–6602

López I, Tournillon A-S, Nylander K, Fåhraeus R (2015) p53-mediated control of gene expression via mRNA translation during Endoplasmic Reticulum stress. Cell Cycle 14: 3373–3378

Love MI, Huber W, Anders S (2014) Moderated estimation of fold change and dispersion for RNA-seq data with DESeq2. Genome Biology 15: 550

Marcel V, Fernandes K, Terrier O, Lane DP, Bourdon JC (2014) Modulation of p53β and p53γ expression by regulating the alternative splicing of TP53 gene modifies cellular response. Cell Death & Differentiation 21: 1377–1387

Marcel V, Perrier S, Aoubala M, Ageorges S, Groves MJ, Diot A, Fernandes K, Tauro S, Bourdon J-C (2010a) Δ160p53 is a novel N-terminal p53 isoform encoded by Δ133p53 transcript. 584: 4463–4468

Marcel V, Vijayakumar V, Fernández-Cuesta L, Hafsi H, Sagne C, Hautefeuille A, Olivier M, Hainaut P (2010b) p53 regulates the transcription of its Δ133p53 isoform through specific response elements contained within the TP53 P2 internal promoter. Oncogene 29: 2691–2700

Markham NR, Zuker M (2008) UNAFold: software for nucleic acid folding and hybridization. Methods Mol Biol 453: 3–31

Mondal AM, Zhou H, Horikawa I, Suprynowicz FA, Li G, Dakic A, Rosenthal B, Ye L, Harris CC, Schlegel R et al (2018) Δ133p53α, a natural p53 isoform, contributes to conditional reprogramming and long-term proliferation of primary epithelial cells. Cell Death & Disease 9: 750

Ohki R, Kawase T, Ohta T, Ichikawa H, Taya Y (2007) Dissecting functional roles of p53 N-terminal transactivation domains by microarray expression analysis. 98: 189–200

Osaki M, Tatebe S, Goto A, Hayashi H, Oshimura M, Ito H (1997) 5-Fluorouracil (5-FU) induced apoptosis in gastric cancer cell lines: role of the p53 gene. Apoptosis 2: 221–226

Pal A, Ghosh PK, Das S (2023) The "LINC" between Δ40p53-miRNA Axis in the Regulation of Cellular Homeostasis. Molecular and cellular biology 43: 335–353

Pal A, Ghosh PK, Ghosh S, Tripathi SK, Patra S, Khan D, Maitra A, Das S (2026) Disruption of the Δ40p53/miR-4671-5p/SGSH axis results in intra-S-phase arrest and poor cancer prognosis. The FEBS journal

Pritchard DM, Potten CS, Hickman JA (1998) The relationships between p53-dependent apoptosis, inhibition of proliferation, and 5-fluorouracil-induced histopathology in murine intestinal epithelia. Cancer research 58: 5453–5465

Ray PS, Grover R, Das S (2006) Two internal ribosome entry sites mediate the translation of p53 isoforms. EMBO Rep 7: 404–410

Robinson MD, McCarthy DJ, Smyth GK (2010) edgeR: a Bioconductor package for differential expression analysis of digital gene expression data. Bioinformatics 26: 139–140

Romar GA, Kupper TS, Divito SJ (2016) Research Techniques Made Simple: Techniques to Assess Cell Proliferation. Journal of Investigative Dermatology 136: e1–e7

Shamloo B, Usluer S (2019) p21 in Cancer Research. Cancers 11

Sharathchandra A, Lal R, Khan D, Das S (2012) Annexin A2 and PSF proteins interact with p53 IRES and regulate translation of p53 mRNA. RNA Biology 9: 1429–1439

Song R, Tian K, Wang W, Wang L (2015) P53 suppresses cell proliferation, metastasis, and angiogenesis of osteosarcoma through inhibition of the PI3K/AKT/mTOR pathway. International journal of surgery (London, England) 20: 80–87

Tomas F, Roux P, Gire V (2024) Interaction of p53 with the Δ133p53α and Δ160p53α isoforms regulates p53 conformation and transcriptional activity. Cell Death & Disease 15: 845

Vilgelm AE, Washington MK, Wei J, Chen H, Prassolov VS, Zaika AI (2010) Interactions of the p53 Protein Family in Cellular Stress Response in Gastrointestinal Tumors. Molecular Cancer Therapeutics 9: 693–705

Vousden KH, Prives C (2009) Blinded by the Light: The Growing Complexity of p53. Cell 137: 413–431

Wang X, Li L, Mok TSK, Tao Q (2018) 8P Noncanonical Wnt11, a tumor suppressive gene by antagonizing canonical Wnt signaling, represents a putative molecularly therapeutic target in lung cancer. Journal of Thoracic Oncology 13: S4–S5

Wu Q, Xu C, Zeng X, Zhang Z, Yang B, Rao Z (2021) Tumor suppressor role of sFRP-4 in hepatocellular carcinoma via the Wnt/β-catenin signaling pathway. Mol Med Rep 23: 336

Xie N, Chen M, Dai R, Zhang Y, Zhao H, Song Z, Zhang L, Li Z, Feng Y, Gao H et al (2017) SRSF1 promotes vascular smooth muscle cell proliferation through a Δ133p53/EGR1/KLF5 pathway. Nature Communications 8: 16016

Yang CM, Kang MK, Jung WJ, Joo JS, Kim YJ, Choi Y, Kim HP (2021) p53 expression confers sensitivity to 5-fluorouracil via distinct chromatin accessibility dynamics in human colorectal cancer. Oncology letters 21: 226

Yee KS, Wilkinson S, James J, Ryan KM, Vousden KH (2009) PUMA- and Bax-induced autophagy contributes to apoptosis. Cell death and differentiation 16: 1135–1145

Zawacka-Pankau JE (2020) The Undervalued Avenue to Reinstate Tumor Suppressor Functionality of the p53 Protein Family for Improved Cancer Therapy-Drug Repurposing. 12: 2717

Zhang X, Groen K, Morten BC, Steffens Reinhardt L, Campbell HG, Braithwaite AW, Bourdon J-C, Avery-Kiejda KA (2022) Effect of p53 and its N-terminally truncated isoform, Δ40p53, on breast cancer migration and invasion. 16: 447-465

Zhao L, Punga T, Sanyal S (2025) Δ133p53α and Δ160p53α isoforms of the tumor suppressor protein p53 exert dominant-negative effect primarily by co-aggregation. eLife 14: RP106469

Zilfou JT, Lowe SW (2009) Tumor suppressive functions of p53. Cold Spring Harbor perspectives in biology 1: a001883

