## Supplementary material for "IRES-mediated translation of Δ160p53 regulates p53 functions and fine-tunes cancer homeostasis": nothing

Supplementary Figure 1:

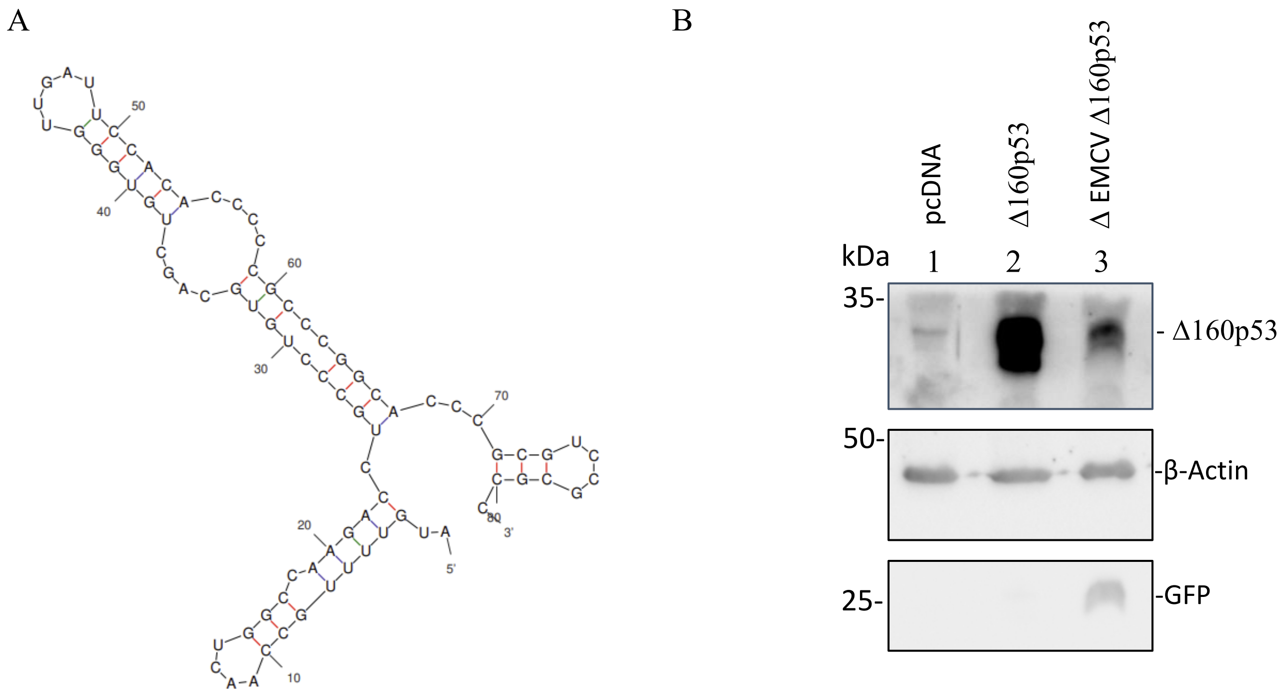


Supplementary Figure 2:

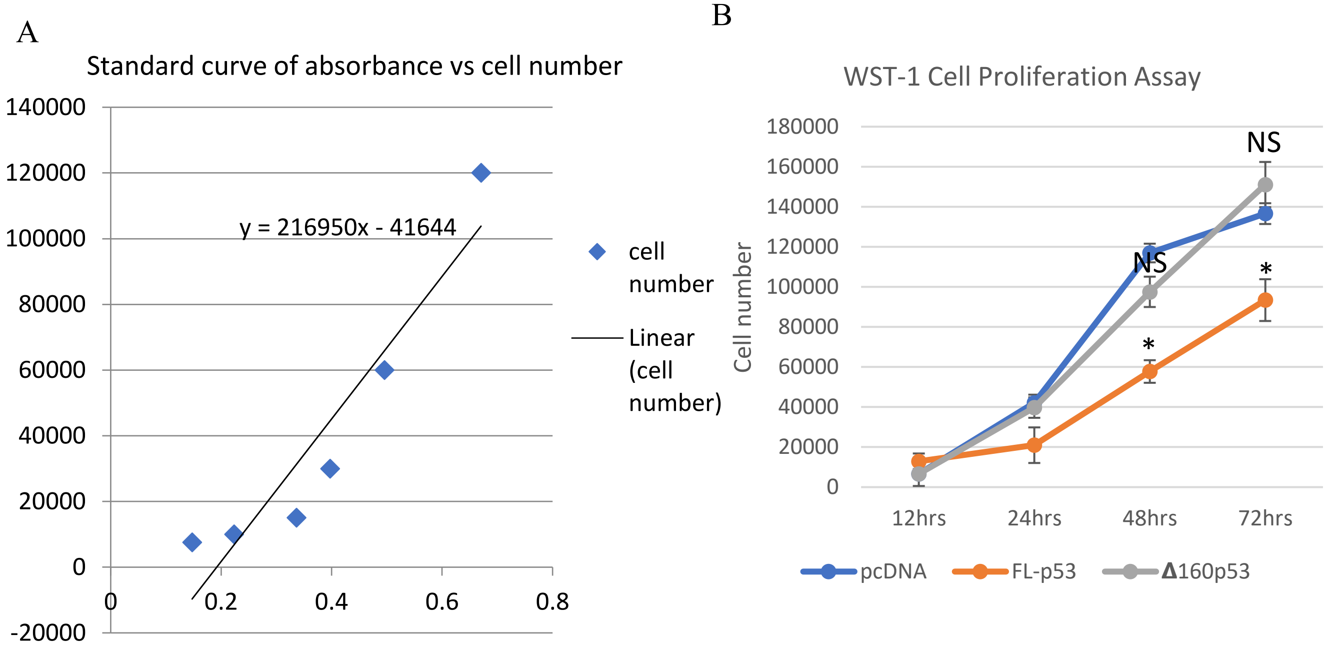

Supplementary Figure 3:

A

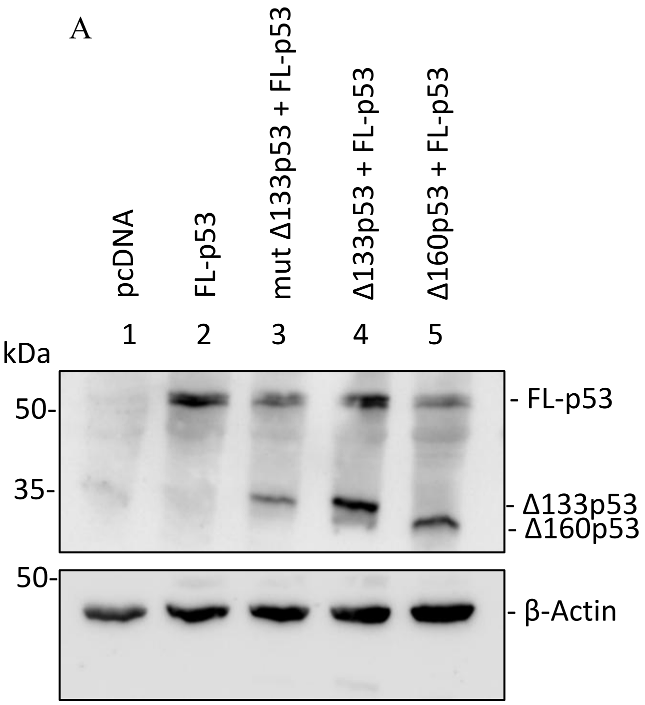


B

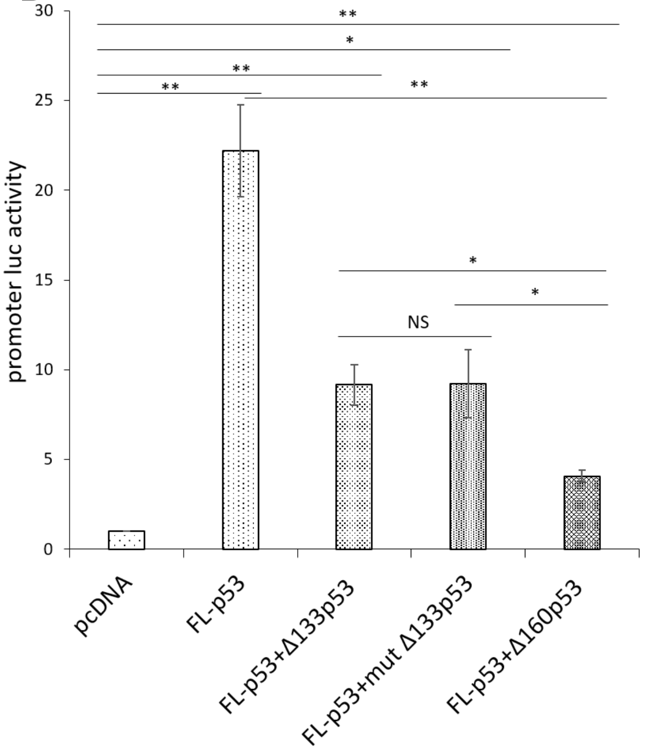


Supplementary Figure 4:

A

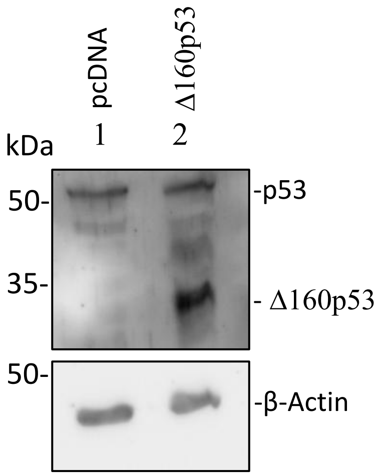


B

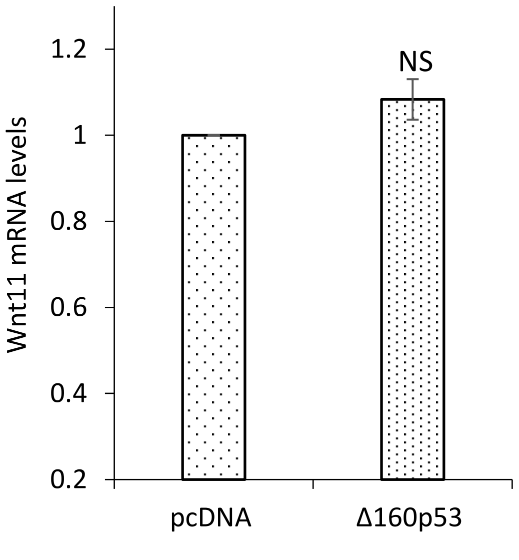

### Supplementary Table 1:

| Pathway Genes | Fold Enrichment | Pathway | Genes |
|---|---|---|---|
| 63 | 12.62135395 | Path:hsa05217 Basal cell carcinoma | TP53 WNT7A WNT7B WNT11 |
| 100 | 11.92717949 | Path:hsa04916 Melanogenesis | ADCY5 EDN1 WNT7A WNT7B WNT11 CAMK2B |
| 69 | 11.52384492 | Path:hsa05031 Amphetamine addiction | ADCY5 GRIN3B ARC CAMK2B |
| 75 | 10.60193732 | Path:hsa04971 Gastric acid secretion | ADCY5 KCNQ1 KCNK10 CAMK2B |
| 96 | 10.35345442 | Path:hsa04713 Circadian entrainment | ADCY5 ADCYAP1 ADCYAP1R1 CAMK2B CACNA1I |
| 85 | 9.354650578 | Path:hsa04911 Insulin secretion | ADCY5 ADCYAP1 ADCYAP1R1 CAMK2B |
| 154 | 9.035742036 | Path:hsa04934 Cushing syndrome | CDKN1A ADCY5 WNT7A WNT7B WNT11 CAMK2B CACNA1I |
| 114 | 8.718698456 | Path:hsa04725 Cholinergic synapse | ADCY5 CHRM4 CHRNA4 KCNQ1 CAMK2B |
| 144 | 8.282763533 | Path:hsa04550 Signaling pathways regulating pluripotency of stem cells | FGFR3 INHBB LHX5 WNT7A WNT7B WNT11 |
| 153 | 7.795542148 | Path:hsa04261 Adrenergic signaling in cardiomyocytes | ADCY5 KCNQ1 ATP2B3 SLC8A3 CAMK2B CACNA2D2 |
| 173 | 6.894323403 | Path:hsa04310 Wnt signaling pathway | SFRP4 TP53 WNT7A WNT7B WNT11 CAMK2B |
| 203 | 6.854700855 | Path:hsa05205 Proteoglycans in cancer | CDKN1A IHH TP53 WNT7A WNT7B WNT11 CAMK2B |
| 148 | 6.715754216 | Path:hsa05224 Breast cancer | CDKN1A TP53 WNT7A WNT7B WNT11 |
| 149 | 6.67068204 | Path:hsa05226 Gastric cancer | CDKN1A TP53 WNT7A WNT7B WNT11 |
| 370 | 6.447124047 | Path:hsa04080 Neuroactive ligand-receptor interaction | CHRM4 CHRNA4 CHRND ADCYAP1 GRIN3B ADCYAP1R1 S1PR3 EDN1 GABRD GRM4 MC5R LYNX1 |
| 157 | 6.330774675 | Path:hsa04390 Hippo signaling pathway | CRB2 GDF6 WNT7A WNT7B WNT11 |
| 252 | 6.310676977 | Path:hsa04020 Calcium signaling pathway | GRIN3B FGFR3 GDNF ATP2B3 RET SLC8A3 CAMK2B CACNA1I |
| 226 | 6.157098555 | Path:hsa04024 cAMP signaling pathway | ADCY5 ADCYAP1 GRIN3B ADCYAP1R1 EDN1 ATP2B3 CAMK2B |
| 300 | 4.638347578 | Path:hsa04010 MAPK signaling pathway | FGFR3 GDNF NGFR RET TP53 CACNA1I CACNA2D2 |
| 529 | 3.757775516 | Path:hsa05200 Pathways in cancer | CDKN1A ADCY5 EDN1 FGFR3 RET TP53 WNT7A WNT7B WNT11 CAMK2B |

### Supplementary Table 2:

| Primer Name | Primer Sequence 5' --> 3' |
|---|---|
| SRPF4 FWD | CTATGACCGTGGCGTGTGCATT |
| SRPF4 REV | GCTTAGGCGTTTACAGTCAACATC |
| CAMK2B FWD | ACACCGTCACTCCTGAAGCCAA |
| CAMK2B REV | GTGCATCATGGATGCTACCGTG |
| WNT7a FWD | AGGAGAAGGCTCACAAATGGGC |
| WNT7a REV | CGGCAATGATGGCGTAGGTGAA |
| ADCY5 FWD | TACACTGCGGTGTCCTTGGTCT |
| ADCY5 REV | GAGTGTAGCCTTGGTGATGTGG |
| WNT11 FWD | CTGTGAAGGACTCGGAACTCGT |
| WNT11 REV | AGCTGTCGCTTCCGTTGGATGT |
| GAPDH Fwd | CAGCCTAGGATCATCAGCAAT |
| GAPDH Rev | GGTCATGAGTCCTTCCACGA |
| Puma fwd | ACCTCAACGCACAGTACGAG |
| Puma rev | CCCATGATGAGATTGTACAGGA |
| p21 fwd | TCTTGTACCCTTGTGCCTCG |
| p21 rev | AGAAGATCAGCCGGCGTTTG |
| Bax fwd | GGCCGGGTTGTCGCCCTTTT |
| Bax rev | CCGCTCCCGGAGGAAGTCCA |
| Luc Fwd | TCCGGATACTGCGATTTTAAG T |
| Luc Rev | CCTGAAGGGATCGTAAAAACAG |
| P3 | CGAAAGTTTATGATCCAGAACAAA |
| P1 | ATCGTTCGTTGAGCGAGTTCTCAA |
| P4 | TTATGTTTTTGGCGTCTTCCAT |
| P2 | TTACACGGCGATCTTTCCGCCC |

### Supplementary Figure legends

**Supplementary Figure-1 Secondary structure of putative IRES and IRES-mediated translation of Δ160p53 in HeLa cells**

(A) Prediction of stable secondary structure of putative internal ribosomal entry site (IRES) using the UNAFOLD algorithm. (B) Western blot analysis of cell extracts from HeLa cells transfected with pcDNA (negative control), Δ160p53 (positive control), and peGFPΔEMCVI3Δ160p53, probed with p53 polyclonal antibody, β-Actin antibody, and green fluorescence protein (GFP) antibody.

**Supplementary Figure-2 WST-1 cell proliferation assay in H1299 cells**

(A) Standard curve of absorbance vs cell number for WST-1 cell proliferation assay in H1299. (B) Cell proliferation represented in a cell number vs time points graph in H1299 stable cells expressing pcDNA, Δ160p53, and FL-p53. Error bars indicate standard deviation (SD). All experiments were performed in three biological replicates ($n = 3$). The criterion for significance was determined using a two-tailed Student's t-test ($*P \leq 0.05$).

**Supplementary Figure-3 Δ160p53-mediated regulation of promoter activation in heterotetrameric conditions in H1299 cells**

(A) Western blot validation of FL-p53, mut Δ133p53 (2nd AUG is mutated, which cannot produce Δ160p53), Δ133p53, Δ160p53 in H1299 transfected with pcDNA, FL-p53, mut Δ133p53, Δ133p53 and Δ160p53 and luciferase constructs (PG13 and pRL-TK), probed with p53 polyclonal antibody, and β-actin antibody. (B) Luciferase assay of H1299 cells after 48 h of transfection with pcDNA, FL-p53, mut Δ133p53, Δ133p53, and Δ160p53, and luciferase constructs (PG13 and pRL-TK). Error bars indicate standard deviation (SD). All experiments were performed in three biological replicates ($n = 3$). The criterion for significance was determined using a two-tailed Student's t-test ($*P \leq 0.05$ or $**P \leq 0.01$).

**Supplementary Figure-4 Δ160p53-mediated regulation of Wnt11 in A549 cells**

(A) Western blot analysis of total proteins isolated from A549 cells transfected with pcDNA, Δ160p53, and luciferase constructs (Wnt11 promoter luc and pRL-TK) probed with p53 polyclonal antibody and β-Actin antibody. (B) Luciferase assay in A549 cells after 48 h of transfection with pcDNA, Δ160p53, and luciferase constructs. All experiments were performed in three biological replicates (n = 3). A two-tailed Student’s t-test determined the criterion for significance.

**Supplementary Table**

**Supplementary Table 1**

Pathway enrichment analysis of significantly upregulated genes using shinyGO 0.82

**Supplementary Table 2**

List of primers used for semiquantitative polymerase chain reaction (PCR), real-time PCR (RT-PCR), and cDNA preparation
